# Inactivation of *Ppp1r15a* minimises weight gain and insulin resistance during caloric excess in mice

**DOI:** 10.1101/313940

**Authors:** Vruti Patel, Guillaume Bidault, Joseph E. Chambers, Stefania Carobbio, Angharad J. T. Everden, Concepción Garcés, Lucy E. Dalton, Fiona M. Gribble, Antonio Vidal-Puig, Stefan J. Marciniak

## Abstract

Phosphorylation of the translation initiation factor eIF2α within the mediobasal hypothalamus is known to suppress food intake, but the role of the eIF2α phosphatases in regulating body weight is poorly understood. Mice deficient in active PPP1R15A, a stress-inducible eIF2α phosphatase, are healthy and more resistant to endoplasmic reticulum stress than wild type controls. We report that when *Ppp1r15a* mutant mice are fed a high fat diet they gain less weight than wild type littermates owing to reduced food intake. This results in healthy leaner *Ppp1r15a* mutant animals with reduced hepatic steatosis and improved insulin sensitivity, albeit with a modest defect in insulin secretion. By contrast, no weight differences are observed between wild type and *Ppp1r15a* deficient mice fed a standard diet. We conclude that mice lacking the C-terminal PP1-binding domain of PPP1R15A show reduced dietary intake and preserved glucose tolerance. Our data indicate that this results in reduced weight gain and protection from diet-induced obesity.

## Introduction

Mechanisms that govern energy balance have evolved to cope with periods of nutrient scarcity interspersed by times of plenty. The intake of excess calories and their storage in adipose tissue is one such mechanism, but it can become maladaptive in the face of a high calorie western diet. The resulting epidemic of obesity is impairing health in part through increasing the prevalence of type 2 diabetes. By targeting the pathways that promote a positive energy balance it may be possible to prevent obesity and reduce its complications.

The endoplasmic reticulum (ER) is an organelle responsible for the biosynthesis of lipids and the folding of secretory proteins [1]. If protein-folding homeostasis (so-called proteostasis) is threatened by the accumulation of misfolded proteins inside the ER, the cell experiences ER stress. Pathways that are triggered by ER stress are collectively called the unfolded protein response (UPR). Because ER stress is frequently observed in tissues that have accumulated excess lipids it is likely that the UPR is also involved in regulating positive energy balance and/or its associated metabolic complications [2].

The UPR comprises three parallel pathways activated by distinct ER stress sensors: IRE1, ATF6 and PERK [3]. PERK plays a critical role in maintaining the health of insulin-secreting beta-cells and so homozygous mutations of the *PERK* gene in Wolcott-Rallison syndrome manifest as early onset insulin-dependent diabetes [4]. In mice, inactivation of the *Perk* gene also causes early beta-cell death [5]. PERK is a member of a kinase family that selectively phosphorylates eIF2α on serine 51 to P-eIF2α to trigger the integrated stress response (ISR) [6, 7]. This phosphorylation of eIF2α inhibits protein synthesis by preventing translation initiation and thus reduces the load of new proteins entering the ER. When eIF2α is phosphorylated, although the synthesis of most proteins is inhibited, the mRNA of the transcription factor ATF4 is preferentially translated [8]. This leads to the induction of PPP1R15A (also known as GADD34), which binds protein phosphatase 1 (PP1) and G-actin to dephosphorylate P-eIF2α [9, 10]. After a delay of several hours this restores normal levels of translation thus enabling the synthesis of targets of the UPR. However, in the context of chronic ER stress the recovery of protein synthesis mediated by PPP1R15A contributes to toxicity [11, 12].

We previously showed that inactivation of the *Ppp1r15a* gene generates phenotypically healthy, fertile mice that have increased resilience to ER stress [12]. Subsequently, loss of PPP1R15A has been shown to protect against tissue damage in a model of ER stress-induced disease [13]. Subtle reduction in the level of P-eIF2α, the substrate of PPP1R15A, in mice heterozygous for the *eIF2*α^*S51A*^ allele causes obesity, hyperleptinaemia and glucose intolerance when the animals are fed a high-fat diet [14, 15]. However, while increased levels of eIF2α phosphorylation within the hypothalamus have been shown to reduce food intake [16], a recent report suggested that PPP1R15A deficient mice, which are impaired in P-eIF2α dephosphorylation, spontaneously become obese [17]. The specific role of PPP1R15A in the regulation of energy balance therefore remains unclear.

We set out to determine the response of PPP1R15A deficient animals to caloric excess using a high-fat diet. Contrary to a previous report [17], we observed *Ppp1r15a^ΔC/ΔC^* mice to gain significantly less weight than wild type littermates when fed a high-fat diet. Metabolic phenotyping revealed reduced food intake in the absence of changes in energy expenditure. Consequently, despite having a subtle defect in insulin secretion, *Ppp1r15a^ΔC/ΔC^* mice were resistant to diet induced obesity, remaining leaner and less insulin resistant than their wild type littermates fed on a high-fat diet.

## Materials and Methods

### Mice

The Guide for the care and use of laboratory animals, Eight edition (2011), was followed. The UK Home Office approved all animal protocols in this study (PPL 80/2491 & PPL 80/2484). Generation of the *Ppp1r15a^ΔC/ΔC^* mouse (originally named *Ppp1r15a^tm1Dron^*), has been described [11]. To minimise effects of genetic background, backcrossing was performed (Speed Congenics, Charles River, UK) until 97% C57BL/6 purity. Mice were fed a standard chow diet (5.1% fat, SAFE diets U8400G10R) or a semisynthetic high-fat diet (60% fat, Research diets D12492i, Research Diets, New Brunswick, NJ). All mice used in this study were female. For treatment with streptozotocin, mice were intraperitoneally injected with 50mg/kg streptozotocin (Sigma□S0130, Dorset, UK) freshly dissolved in sodium citrate pH 4.5.

### Metabolic profiling

We performed glucose tolerance tests after a 16□hour fast (5 p.m. to 8 a.m.) and insulin tolerance tests after a 4□hour fast (8 a.m. to 12 p.m.). Body composition was measured by time domain NMR (Bruker Optics). Daily food intake measurements were carried out on singly-housed mice in surgery bedding lined cages. Energy expenditure was measured by indirect-calorimetry in a Metatrace system (Creative Scientific, UK) at 21°C attached to a custom□built oxygen and carbon dioxide monitoring system (Minimox system built by P. Murgatroyd). Carbon dioxide concentration in room air and air leaving each cage were measured every 11 minutes. Activity was assessed by beam breaks set 2.5 cm apart. Activity measurements were taken to be total beam breaks, measured every 8 minutes.

### Glucose and insulin tolerance tests

Overnight fasted mice were injected intraperitoneally with 1g/kg glucose, calculated using the average bodyweight of the cohort, and blood glucose was measured for 2 hours. For the insulin tolerance tests, mice were fasted for 4 hours, basal glucose measured, and then mice were injected intraperitoneally with 0.75U/kg of insulin, calculated using the average bodyweight of the *Ppp1r15a^ΔC/ΔC^* cohort. Measurement of plasma insulin and free fatty acid was carried out using the Meso Scale Discovery (MSD) assays at the Core Biochemical Assay Laboratory (CBAL), Cambridge, UK.

### Histology

Primary antibodies used were: guinea-pig anti-insulin (ab7842, 1:200; Abcam), rabbit anti-glucagon (SC13091, 1:200; SantaCruz, Santa Cruz, CA, USA). Sections were counterstained using Hoechst (1:1300, Invitrogen, UK). Stained sections were imaged using either the incubated Zeiss LSM510 META Confocal Microscope, Zeiss LSM710 META Confocal Microscope, Zeiss AxioImager Motorised Upright Microscope. Images were analysed using Zen software.

### Cells and reagents

Wild type and *Ppp1r15a^ΔC/ΔC^* MEFs were cultured as described [12]. Prior to experimentation, palmitate was first conjugated to BSA. Palmitic acid (Sigma□P0500, Dorset, UK) was prepared to a concentration of 100mM in 100% (v/v) ethanol and heated at 60°C. This was added to media supplemented with 5% (w/v) fatty□acid free, low endotoxin BSA (Sigma□A8806, Dorset, UK), to prepare a 10X stock and sonicated to ensure conjugation and solubilisation.

### Immunoblots

Cell lysates were prepared and resolved as described [12]. Primary antibodies used were: rabbit anti-PPP1R15A (10449-1-AP, 1:1000; Proteintech, Manchester, UK), rabbit anti-phospho-eIF2α (3597, 1:1000; Cell Signaling), mouse anti-actin (ab3280, 1:1000; Abcam), rabbit anti-ATF4 (C-20, 1:500; SantaCruz, Santa Cruz, CA, USA), mouse anti-total eIF2α (AHO0802, 1:1000; Invitrogen, Thermo Fisher Scientific, Waltham, MA, USA); guinea pig anti-insulin (ab7842, 1:1000; Abcam), rabbit anti-GAPDH (14C10) (2118, 1:1000; Cell Signaling), rabbit anti-UCP1 (ab10983, 1:1000; Abcam), rabbit anti-CHOP (1:1000; kind gift from Professor David Ron, Cambridge, UK), rabbit anti-alpha/beta tubulin (2148, 1:1000; Cell Signalling, Hitchin, UK).

### RNA extraction and real-time PCR

Total RNA was extracted from mammalian cells using the Qiagen RNeasy Mini Kit (Qiagen Ltd., UK), and from animal tissue using TRIzol reagent (15596026, ThermoFisher). cDNA was reverse transcribed with M□MLV Reverse Transcriptase (Invitrogen, UK), using an Oligo-(dt)_15_ primer (Promega, UK). Real□time PCR was performed using an ABI7900-HT-Fast device (Applied Biosystems, Carlsbad, UK). Relative fold□changes of expression were determined using the relative standard curve method or, were stated in the legend, BestKeeper analysis [18]. Primer sequences used can be found in Supplementary Table S1.

### Statistics

Data were analysed using unpaired Student’s t□tests or analysis of variance (ANOVA) with Bonferroni *post hoc* correction (Prism, Graphpad, USA).

## Results

### Inactivation of PPP1R15A reduces ISR signalling and protects from lipotoxicity *in vitro*

As expected, when mouse embryonic fibroblasts (MEFs) generated from wild type embryos were treated with the ER stressor thapsigargin, full length PPP1R15A protein was induced progressively over an 8-hour period (Fig 1a & 1b). This mirrored the transcriptional induction of *Ppp1r15a* mRNA (Fig 1c & 1d). In *Ppp1r15a^ΔC/ΔC^* MEFs, PPP1R15A-ΔC was induced over a similar time course, but to significantly lower levels. A primer pair targeting the deleted exon 3, served as a convenient control for genotype (Fig 1c); while, primers within the remaining *Ppp1r15a^ΔC^* allele enabled us to use endogenous *Ppp1r15a* mRNA as a readout of the P-eIF2α-ATF4 (ISR) pathway. *Ppp1r15a^ΔC/ΔC^* showed no induction of exon 3, but endogenous *Ppp1r15a^ΔC^* was induced albeit to a much lower level than the wild type allele (Fig 1d). Consistent with this, P-eIF2α peaked in wild type cells at 0.5 hours after treatment with thapsigargin, falling back to baseline by 8 hours as PPP1R15A levels rose, while P-eIF2α rose monotonically in the *Ppp1r15a^ΔC/ΔC^* MEFs (Fig 1a & 1b). This confirmed that *Ppp1r15a^ΔC/ΔC^* cells are impaired in the dephosphorylation of P-eIF2α during ER stress.

**Figure 1.**
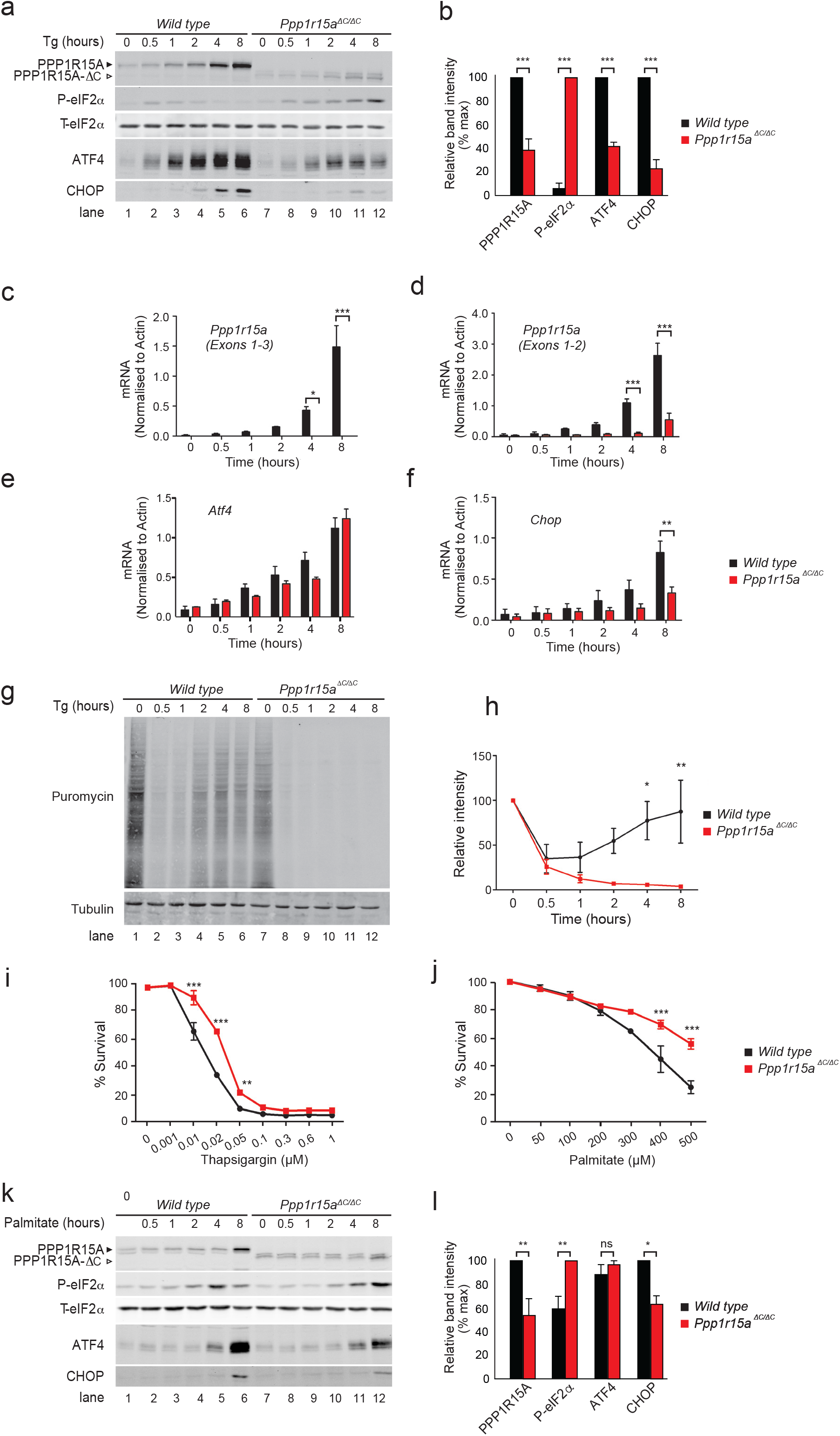
Inactivation of PPP1R15A reduces ER stress and protects against lipotoxicity *in vitro*. (a) Immunoblot for PPP1R15A, phosphorylated eIF2α (P-eIF2α), total eIF2α (T-eIF2α), ATF4 and CHOP in lysates of wild type and *Ppp1r15a^ΔC/ΔC^* mouse embryonic fibroblasts (MEFs) following treatment with thapsigargin (Tg) 300nM for indicated times. Proteins of the expected sizes are marked with a solid triangle for PPP1R15A or an open triangle for PPP1R15A-ΔC. (b) Quantification of (a) using ImageJ software. N=3; mean ± SEM; P value calculated by two-way ANOVA. (c-f) Wild type and *Ppp1r15a^ΔC/ΔC^* MEFs were treated with thapsigargin 300nM for indicated times and RNA was prepared. *Ppp1r15a (Exons 1-3), Ppp1r15a (Exons 1-2), Atf4, Chop* mRNA were quantified relative to *beta-actin* by qRT-PCR. N=3; mean ± SEM; P value calculated by two-way ANOVA. (g) Immnoblot for puromycin (indicating rate of translation) and tubulin in lysates of wild type or *Ppp1r15a^ΔC/ΔC^* MEFs following treatment with thapsigargin (Tg) 300nM for indicated times. Ten minutes prior to harvesting, puromycin was added to the culture medium at a final concentration of 10ng/mL. (h) Immunoreactivity to puromycin within lysates served as a marker of protein translation and was quantified using ImageJ software. N=3; mean ± SEM; P value calculated by two-way ANOVA with Bonferroni *post hoc* test. (i) MTT assays were carried out to measure cell viability of *wild type* or *Ppp1r15a^ΔC/ΔC^* MEFs following treatment with the indicated concentrations of thapsigargin for 48 hours. N=3; mean ± SEM; P value calculated by two-way ANOVA with Bonferroni *post hoc* test (j) MTT assays were carried out to measure cell viability of wild type or *Ppp1r15a^ΔC/ΔC^* MEFs following treatment with the indicated concentrations of palmitate for 48 hours. N=3; mean ± SEM; P value calculated by two-way ANOVA with Bonferroni *post hoc* test (k) Immunoblot for PPP1R15A, P-eIF2α, T-eIF2α, ATF4 and CHOP in lysates of wild type and *Ppp1r15a^ΔC/ΔC^* mouse embryonic fibroblasts (MEFs) following treatment with palmitate 400μm for indicated times. Proteins of the expected sizes are marked with a solid triangle for PPP1R15A or an open triangle for PPP1R15A-ΔC. (l) Quantification of (A) using ImageJ software. N=3; mean ± SEM; P value calculated by two-way ANOVA. *** p<0.001, ** p<0.01, * p<0.05.

The induction of ATF4 protein was significantly reduced in *Ppp1r15a^ΔC/ΔC^* MEFs compared with wild type controls despite similar levels of *Atf4* mRNA (Fig 1a, 1b & 1e). This reflected a more intense inhibition of translation in *Ppp1r15a^ΔC/ΔC^* MEFs (Fig 1g & 1h). Consistent with this, cells treated with tunicamycin, which induces ER stress more slowly and so inhibits translation more weakly, showed no deficit of ATF4 in the *Ppp1r15a^ΔC/ΔC^* MEFs (Supplementary Fig 1). The lower levels of ATF4 in thapsigargin-treated *Ppp1r15a^ΔC/ΔC^* cells reduced induction of its target gene *Chop* (Fig 1f) and so contributed to the reduced induction of endogenous *Ppp1r15a^ΔC^* mRNA (Fig 1d).

The reduction in ISR signalling in thapsigargin-treated *Ppp1r15a^ΔC/ΔC^* cells correlated with reduced toxicity of this agent. When wild type and *Ppp1r15a^ΔC/ΔC^* MEFs were treated for 48 hours with thapsigargin, PPP1R15A-deficient cells required significantly higher concentration of drug to reduce viability (Fig 1i). The saturated free fatty acid palmitate causes cell death in part via the induction of ER stress [19-22] and, like thapsigargin, this may involve the depletion of ER calcium [19, 23]. We therefore tested whether inactivation of PPP1R15A would protect cells from the lipotoxicity of palmitate. Once again, the PPP1R15A-deficient cells required significantly higher levels of palmitate to reduce viability (Fig 1j). Therefore, we concluded that deficiency of PPP1R15A reduced the lipotoxic insult induced by palmitate *in vitro*. As had been observed for thapsigargin, palmitate caused significantly higher levels of P-eIF2α and lower levels of CHOP in *Ppp1r15a^ΔC/ΔC^* MEFs than in controls (Fig 1k & 1l).

### Inactivation of PPP1R15A reduces weight gain of mice fed a high-fat diet and prevents diet-induced hepatic steatosis

Reduced phosphorylation of eIF2α has been shown to enhance weight gain of mice fed a high-fat diet [14]. However, loss of the eIF2α phosphatase PPP1R15A has also been found to cause obesity in male mice, but not in females [17, 24]. To examine the effect of PPP1R15A expression in females in greater detail, we randomly assigned 6-week old female wild type and *Ppp1r15a^ΔC/ΔC^* littermates to normal chow or 60% high-fat diet. Bodyweights were monitored weekly for the following 10 weeks (Fig 2a). Strikingly, only when fed a 60% high-fat diet, *Ppp1r15a^ΔC/ΔC^* animals gained less weight than their wild type littermates (Fig 2a & 2b). This reduced weight did not arise from a runty phenotype, as body length was not significantly different between the two genotypes (Fig 2c).

**Figure 2.**
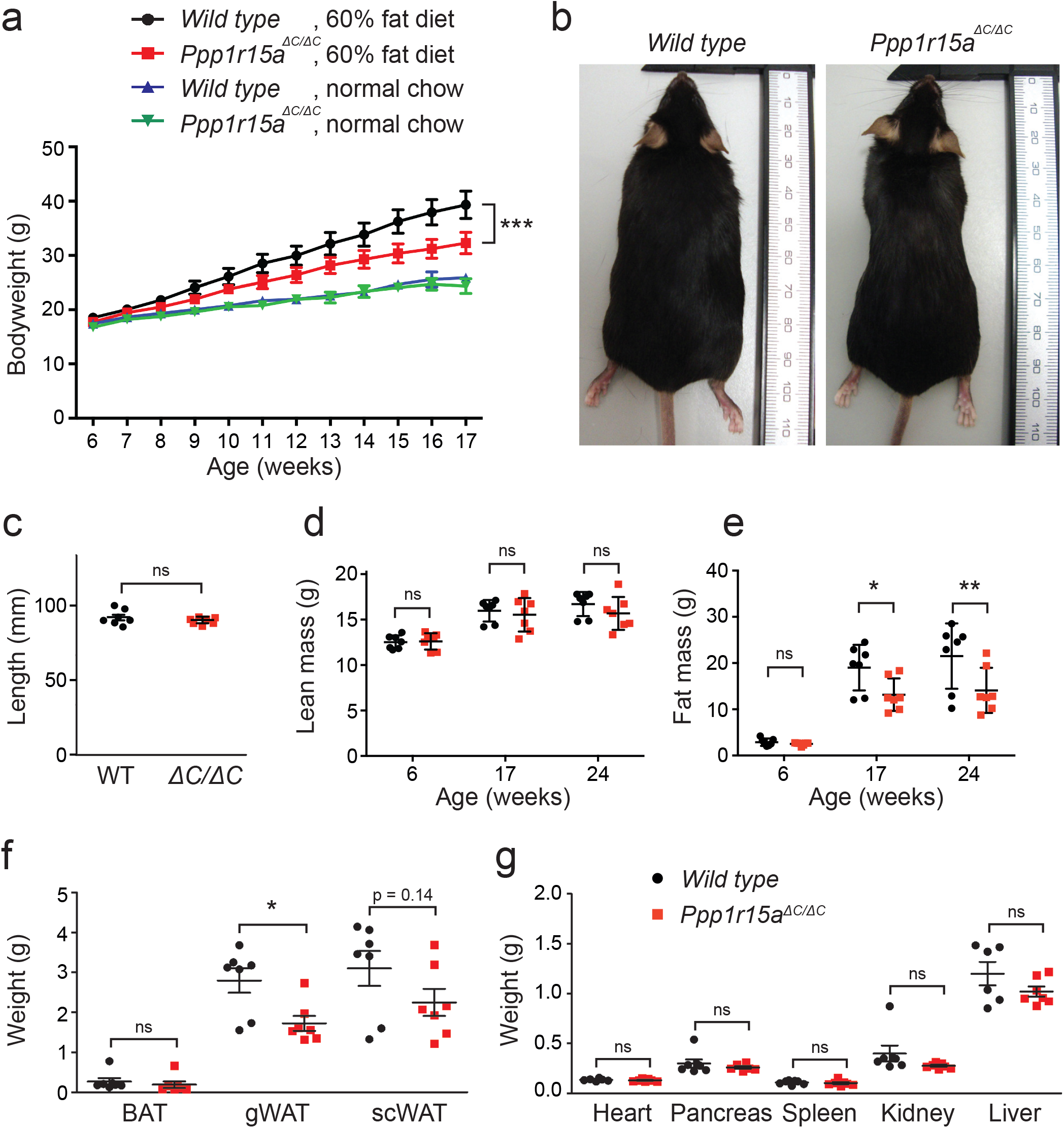
Inactivation of PPP1R15A reduces weight gain following a 60% high-fat diet *in vivo*. (a) At 6 weeks of age, wild type and *Ppp1r15a^ΔC/ΔC^* mice were fed either standard chow diet or 60% high-fat diet and their bodyweights measured. P value calculated by two-way ANOVA. (b) Representative images of wild type and *Ppp1r15a^ΔC/ΔC^* mice fed a 60% high-fat diet for 18 weeks. (c) Length of each mouse was measured at time of harvest. (d-e) Using time domain nuclear magnetic resonance (TD-NMR), the (d) lean mass, and (e) fat mass of the mice were measured prior to high-fat feeding (6-weeks), during high-fat feeding (17-weeks) and at the end of the study (24-weeks). (f) The wet weights of each adipose tissue were measured at harvest. Brown adipose tissue (BAT); gonadal white adipose tissue (gWAT) and subcutaneous white adipose tissue (scWAT). (g) The organs of each mouse were also harvested at the end of the study and wet weights measured. N=7; mean plotted ± SEM; unpaired Student’s unpaired t-test used for statistical analysis. *** p<0.001, ** p<0.01, * p<0.05.

We then used time domain□NMR to investigate whether the body weight differences reflected differences in fat or lean mass. Lean mass was not different between the two genotypes at 6, 17 or 24 weeks (Fig 2d). However, although accretion of fat mass rose progressively in both groups, the increase was significantly attenuated in the *Ppp1r15a^ΔC/ΔC^* animals at 17 and 24 weeks (Fig 2e). Organ weight measurements at 24 weeks were not different between genotypes – only gonadal white adipose tissue mass was significantly reduced in the *Ppp1r15a^ΔC/ΔC^* animals compared with the wild type group (Fig 2f & 2g). The 1g difference in the weight of subcutaneous white fat (Fig 2f) could not account entirely for the difference in mean weight between the two groups (8g at 17 weeks of age; Fig 2a & 2b). Since the time domain□NMR showed this difference arose from fat rather than lean mass, it is likely that non-measured sources of fat, e.g. within the periteum, also contributed to the reduced corpulence of the *Ppp1r15a^ΔC/ΔC^* animals compared with their sisters.

These findings confirmed that differences in bodyweight between the genotypes resulted from reduced fat expansion in the *Ppp1r15a^ΔC/ΔC^* mice rather than differences in linear growth *per se*.

In accordance with a healthy lean phenotype observed in *Ppp1r15a^ΔC/ΔC^* mice, histological analysis and neutral lipid staining revealed reduced hepatic steatosis *in Ppp1r15a*Δ*C/*Δ*C* mice fed a high-fat diet compared to the wild type controls (Fig 3a). It has previously been described that over-expression of PPP1R15A attenuates hepatic lipid biogenesis by preventing the accumulation of P-eIF2α and thus skewing lipogenic transcriptional programmes [25]. However, our analysis of the expression of genes involved in hepatic lipid metabolism in wild type and *Ppp1r15a^ΔC/ΔC^* mice fed on high-fat diet for 12 weeks revealed no differences between the two genotypes in *Pparg, Scd1, Pck1, Fasn* and *Acaca* (Fig 3b-f). Furthermore, expression of *Cebpb*, a positive regulator of hepatic Ppar□ [26, 27], was similar in both genotypes (Fig 3g). These results suggest that the reduced hepatic steatosis of *Ppp1r15a^ΔC/ΔC^* mice was related to a healthy phenotype resulting from decreased exogenous input and reduced adiposity.

**Figure 3.**
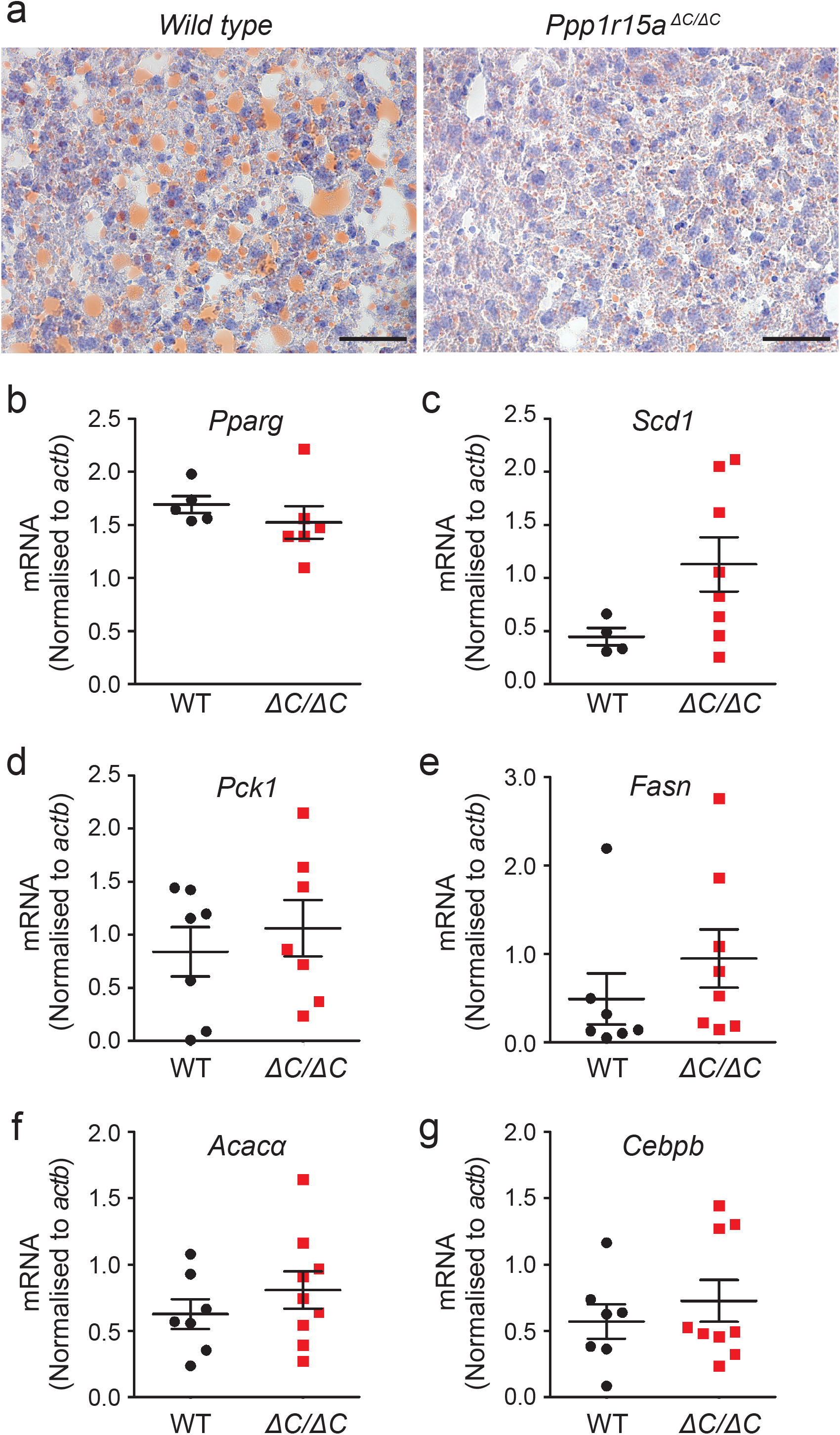
*Ppp1r15a^ΔC/ΔC^* mice show reduced hepatic steatosis following high-fat feeding but no changes in expression of hepatic lipogenic enzymes. (a) To determine hepatic liver content, liver dissected from wild type and *Ppp1r15a^ΔC/ΔC^* mice was frozen in liquid nitrogen. Fifteen-micrometre sections of liver were cut using a cryostat and stained using Oil Red O lipid stain and counterstained with haematoxylin (OROH). Representative images are shown. Scale bar: 50μm. (b-g) RNA was prepared from liver taken from *wild type* (WT) and *Ppp1r15a^ΔC/ΔC^* (ΔC/ΔC) mice fed a high-fat diet for 12 weeks. *Pparg, Scd1, Pck1, Fasn, Acaca*, and *Cebpb* mRNA were quantified relative to *actb* by qRT-PCR. N=4-9; mean plotted ± SEM; none of the mRNA levels were significantly different by unpaired Student’s unpaired t-test.

### Ppp1r15a mutant animals consume less high-fat food without altering energy expenditure

We then investigated the factors contributing to the resistance of *Ppp1r15a^ΔC/ΔC^* mice to high-fat diet-induced obesity. As shown in Fig 2, by 12 weeks on a high-fat diet the bodyweights of wild type and *Ppp1r15a^ΔC/ΔC^* mice were diverging. Therefore at 12 weeks, food intake of singly-housed mice was measured over a 2-week period. This was followed by metabolic rate measurements using indirect calorimetry for a further 48 hours. Food intake, fat mass gained and energy expenditure were calculated as total energy per day for each mouse (Fig 4a). *Ppp1r15a^ΔC/ΔC^* mice showed a significant 16% reduction in food intake compared with wild type controls indicating that inactivation of PPP1R15A selectively decreased the consumption of the high-fat diet (Fig 4a). In the absence of a conditional allele of *Ppp1r15a* it remains to be determined if these differences stem from a function of PPP1R15A in central or peripheral tissues. With respect to energy dissipation, we observed no significant differences in energy expenditure between the two genotypes, nor differences in the expression of the thermoregulatory genes, *Pgc1α*, *Elovl3* and *Dio2,* either in brown (Fig 4b-d) or white adipose tissue (Fig 4e-g). In the brown adipose tissue, we observed a small increase in *Ucp1* mRNA and protein (Fig 4h-k), which was not associated with a decreased respiratory exchange ratio (RER) or an increase in energy expenditure, indicating no preferential oxidation of fatty acids in the *Ppp1r15a^ΔC/ΔC^* mice (Fig 4l). These results argue against PPP1R15A having a major impact on the thermogenic activity of brown and subcutaneous white adipose tissue. Additionally, physical activity rates were indistinguishable between the genotypes (Fig 4m). Thus, the main contributor to resistance to weight gain was decreased food consumption.

**Figure 4.**
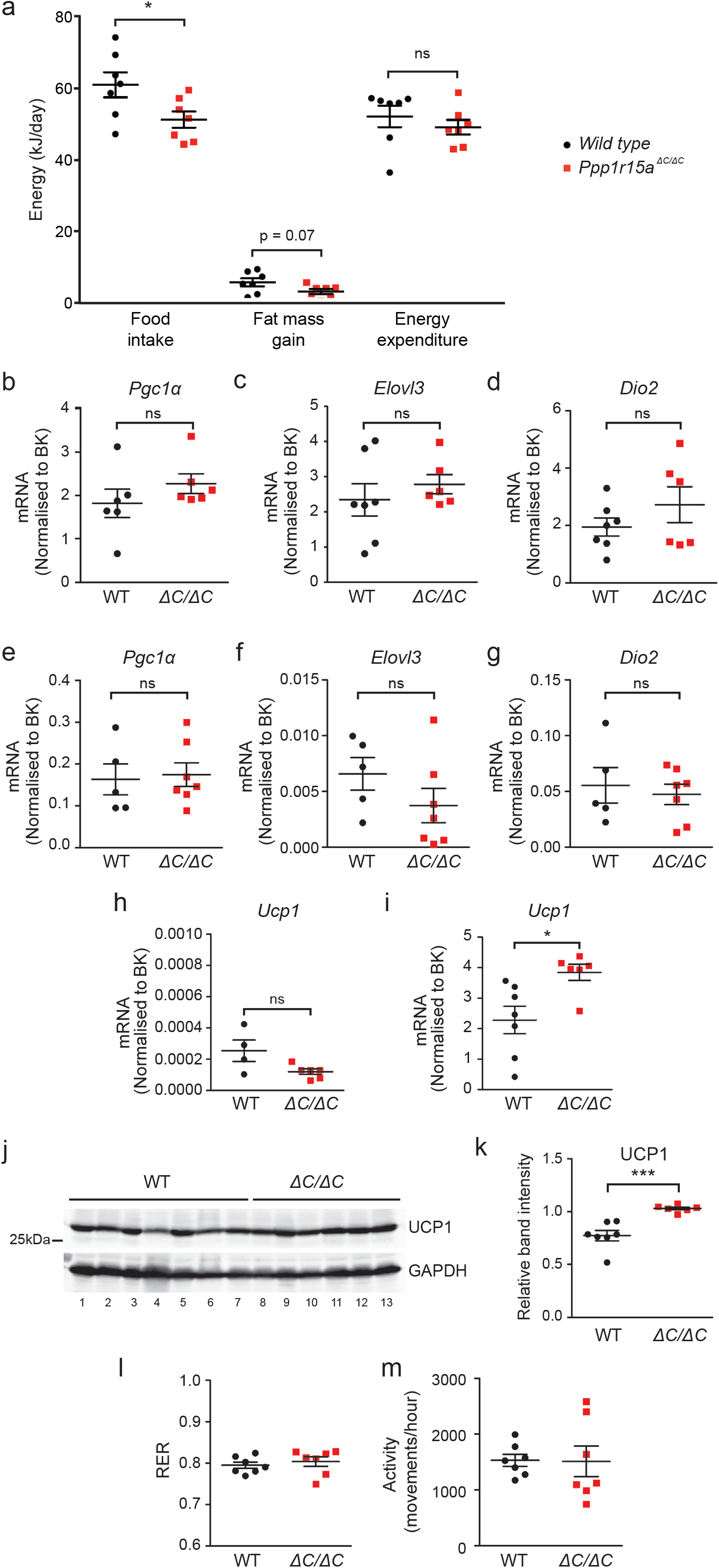
Metabolic phenotyping of *wild type* and *Ppp1r15a^ΔC/ΔC^* mice indicate reduced food intake following high-fat feeding. (a) Mice were single housed at 17-weeks old and food intake was measured three times a week, for 2-weeks. Following food intake measurements, indirect calorimetry was carried out over 48 hours. Food intake, fat mass gain and energy expenditure were calculated. RNA was prepared from brown adipose tissue taken from wild type (WT) and *Ppp1r15a^ΔC/ΔC^* (ΔC/ΔC) mice fed a high-fat diet for 18 weeks. P value calculated by unpaired Student’s t-test. (b-d) *Pgc1a, Elovl3* and *Dio2* mRNA were quantified relative to the geometrical mean of *actb, 36b4, 18S* and *B2M* by qRT-PCR referred to as BestKeeper (BK). RNA was prepared from subcutaneous brown adipose tissue taken from wild type and *Ppp1r15a^ΔC/ΔC^* mice fed a high-fat diet for 18 weeks. (e-g) *Pgc1a, Elovl3* and *Dio2* mRNA were quantified relative to the geometrical mean of *actb, 36b4, 18S* and *B2M* by qRT-PCR referred to as BK analysis. RNA was prepared from white adipose tissue taken from wild type and *Ppp1r15a^ΔC/ΔC^* mice fed a high-fat diet for 18 weeks. (h) RNA was prepared from subcutaneous white adipose tissue taken from wild type and *Ppp1r15a^ΔC/ΔC^* mice fed a high-fat diet for 18 weeks and *Ucp1* mRNA was quantified relative to *actb, 36b4, 18S* and *B2M* by qRT-PCR referred to as BK. (i) RNA was prepared from brown adipose tissue taken from wild type and *Ppp1r15a^ΔC/ΔC^* mice fed a high-fat diet for 18 weeks and *Ucp1* mRNA expression was quantified relative to *actb, 36b4, 18S* and *B2M* by qRT-PCR and BK analysis. (j) Immunoblot for UCP1 and GAPDH of tissue lysates prepared from brown adipose tissue harvested from wild type and *Ppp1r15a^ΔC/ΔC^* mice fed a high-fat diet for 18 weeks. (k) Quantification of (D) using ImageJ software. N=4-7, mean band intensity plotted ± SEM; unpaired Student’s t-test used for statistical analysis. (l-m) Indirect calorimetry was also used to measure the (l) respiratory exchange ratio and (m) activity of the mice over 48 hours. N=7; mean plotted ± SEM; unpaired Student’s t-test used for statistical analysis. *** p<0.001, * p<0.05.

### Glucose tolerance and insulin tolerance testing in wild type and *Ppp1r15a^ΔC/ΔC^* mice

Reduced body weight is not always a sign of health as it is observed in animals with primary adipose tissue dysfunction or lipodystrophy. Thus it was important to determine if the differences in weight gain were associated with improved glucose homeostasis in *Ppp1r15a^ΔC/ΔC^* mice. The glucose tolerance test of mice fed a high-fat diet unexpectedly revealed a hyperglycaemic response in *Ppp1r15a^ΔC/ΔC^* animals at 20 and 30 minutes following the intraperitoneal injection of glucose (Fig 5a). This suggested that despite being leaner, the *Ppp1r15a^ΔC/ΔC^* mice were slightly less glucose tolerant than their wild type littermates. This counterintuitive effect was relatively modest, as the areas under the blood glucose curves were not significantly different between wild type and *Ppp1r15a^ΔC/ΔC^* mice (Fig 5b). Thus, this could have been the result of insulin resistance or impaired insulin secretion by the *Ppp1r15a^ΔC/ΔC^* mice, although basal and 30 minute insulin levels after glucose administration were not measurably different (Fig 5c).

**Figure 5.**
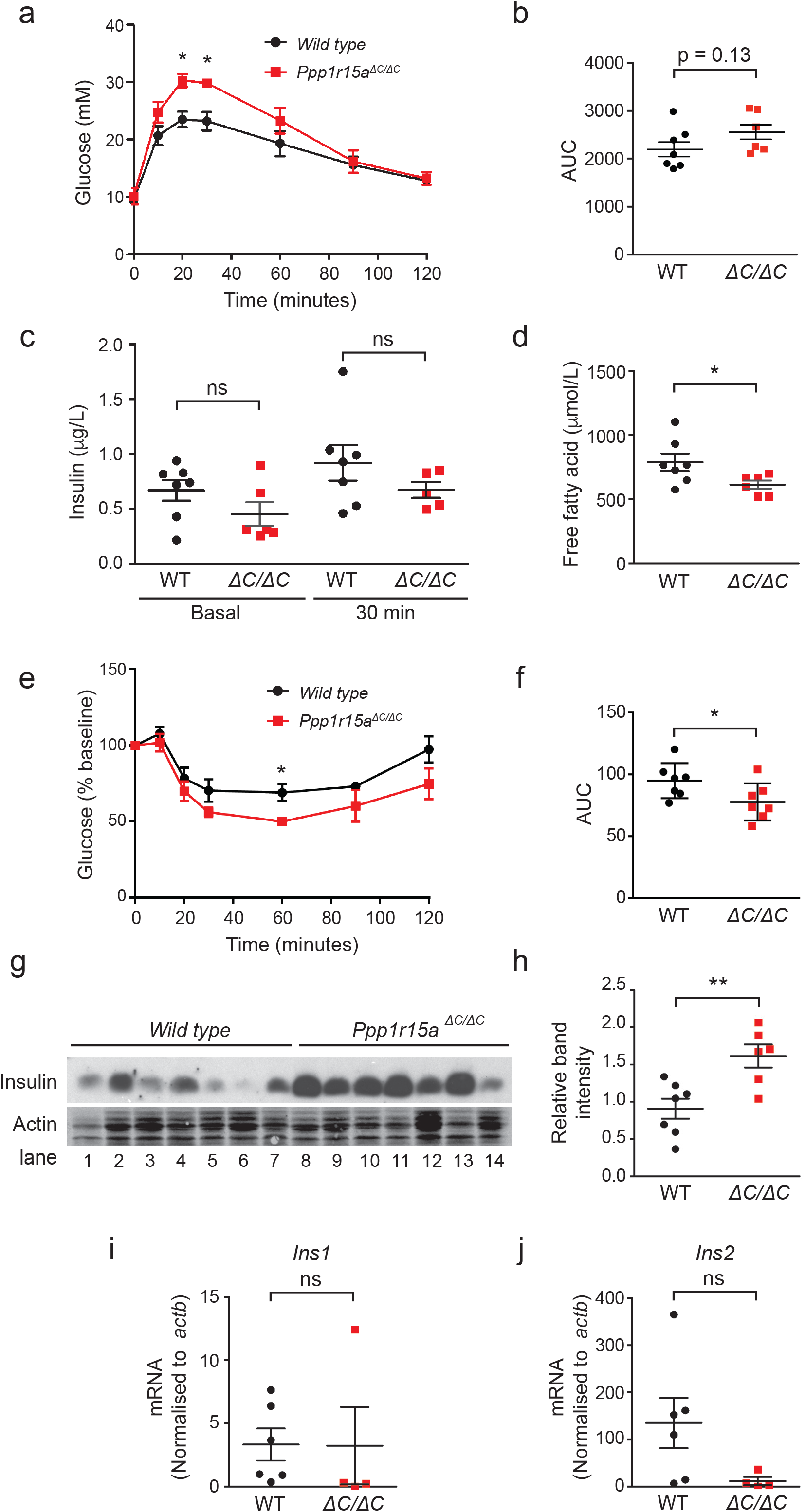
Glucose tolerance and insulin tolerance testing of wild type and *Ppp1r15a^ΔC/ΔC^*mice. (a) At age 22 weeks (16 weeks on high-fat diet), the mice were fasted for 16 hours, after which mice were intraperitoneally injected with 1g/kg glucose, where the dose used was calculated using the average bodyweight of the full cohort of mice. Glucose levels were then measured at 10, 20, 60, 90 and 120 minutes post-injection. P value was calculated by two-way ANOVA with post hoc Bonferroni test. (b) Quantification of the glucose tolerance test (GTT) carried out using the area under the curve (AUC). (c) Blood samples were collected at 0 minutes (basal), prior to glucose injection and at 30 minutes post-injection. Serum was separated out and measured for insulin. P value calculated by unpaired Student’s t-test. (d) Free fatty acid levels in the basal serum sample were also measured. P value calculated by unpaired Student’s t-test. (e) At age 23 weeks (17 weeks on high-fat diet), the mice were fasted for 4 hours, after which mice were intraperitoneally injected with 0.75U/kg insulin, where the dose used was calculated using the average bodyweight of the full cohort of mice. Glucose levels were then measured at 10, 20, 60, 90 and 120 minutes post-injection. P value was calculated by two-way ANOVA with post hoc Bonferroni test. (f) Quantification of the insulin tolerance test (ITT) was carried out using the AUC. P value calculated by unpaired Student’s t-test. (g) Immunoblot for insulin and actin in protein lysates prepared from whole pancreas extracted from wild type and *Ppp1r15a^ΔC/ΔC^* mice fed a high-fat diet for 18 weeks. (h) Quantification of (g) using ImageJ software. P value calculated by unpaired Student’s t-test. (i-j) Total RNA was extracted from whole pancreas of wild type and *Ppp1r15a^ΔC/ΔC^* mice fed a high-fat diet for 18 weeks. *Ins1* and *Ins2* mRNA were quantified relative to *actb* by qRT-PCR. N=4-7; mean plotted ± SEM; unpaired Student’s unpaired t-test used for statistical analysis.

To determine their insulin sensitivity, we performed an insulin tolerance test on animals fed a high-fat diet. Following the injection of 0.75U/kg of insulin, blood glucose fell in both groups, although to a significantly greater degree in the *Ppp1r15a^ΔC/ΔC^* mice (Fig 5e-f). This was seen as both a lower nadir in glucose level and a significantly different area under the blood glucose curve. In line with their leaner phenotype, basal levels of circulating free fatty acids in high-fat diet-fed *Ppp1r15a^ΔC/ΔC^* mice were also significantly lower compared to wild type controls (Fig 5d). This indicated that, in keeping with their leanness and reduced free fatty acid levels, *Ppp1r15a^ΔC/ΔC^* animals were more insulin sensitive than their wild type controls. This further suggested a defect in insulin secretion since *Ppp1r15a^ΔC/ΔC^* mice were less glucose tolerant than their wild type littermates.

Of note, although the basal levels of fasting circulating insulin were not significantly different between the two genotypes (Fig 5c), western blot analysis revealed higher levels of insulin within the pancreata of unfasted *Ppp1r15a^ΔC/ΔC^* animals compared with wild type controls (Fig 5g-h) despite similar levels of insulin mRNA (Fig 5i-j). This also suggested a defect in the secretion of insulin. Loss of PPP1R15A therefore caused opposing effects on glucose homeostasis: on the one hand improving insulin sensitivity, most likely via a negative energy balance, while also impairing early phase insulin secretion allowing higher peak glucose levels following glucose challenge.

### Inactivation of PPP1R15A improves glucose tolerance in a model of beta-cell exhaustion

To determine if inactivation of PPP1R15A would have a net beneficial or detrimental effect in metabolic disease, we set out to model the later stages of type 2 diabetes wherein beta-cell mass falls in the face of sustained insulin resistance [28]. To achieve this, we fed mice a high-fat diet then treated with the beta-cell toxin streptozotocin at a dose optimized to cause subtotal beta-cell loss. At age 6 weeks, mice were placed on 60% high-fat diet and at 12 weeks were injected with 50mg/kg streptozotocin on five consecutive days to mimic the loss of beta-cell mass seen in advanced type 2 diabetes [28]. As expected, streptozotocin injection induced weight loss (Fig 6a) and a progressive increase in fasting glucose in both groups (Fig 6b). Immunofluorescence staining confirmed that islet morphology appeared similar between the genotypes prior to injection with streptozotocin, and both genotypes responded with similar reductions in islet size and falls in circulating insulin (Fig 6c-d).

**Figure 6.**
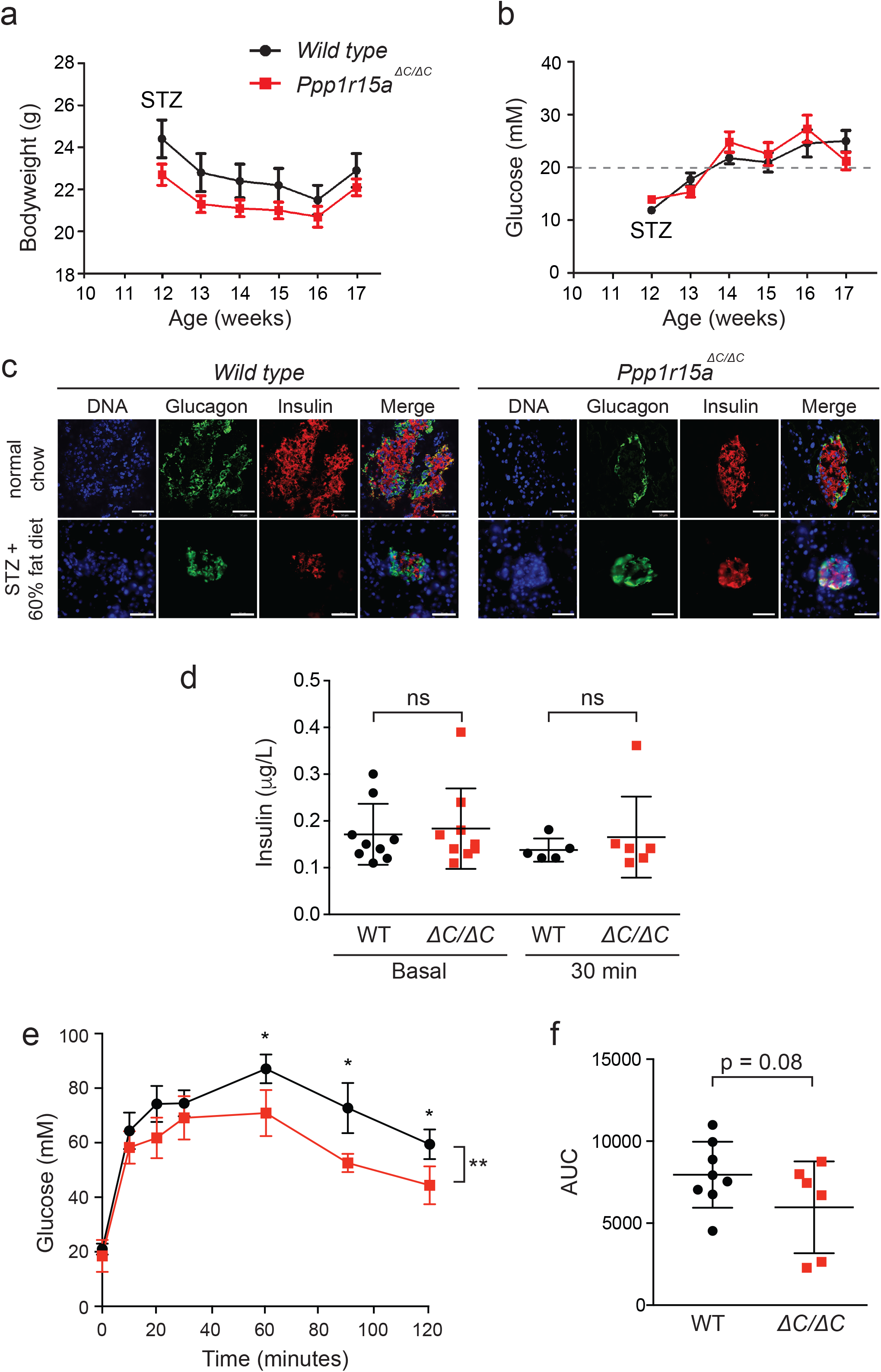
Induction of diabetes and measurement of pancreas function following high-fat feeding and streptozotocin treatment in wild type and *Ppp1r15a^ΔC/ΔC^*mice. (a) Six-week old mice were put on a 60% high-fat diet for six weeks. At age 12 weeks, mice were injected with 50mg/kg streptozotocin (STZ) on 5 consecutive days. Bodyweights of the mice were recorded for 5 weeks post STZ treatment. (b) Non-fasted blood glucose of the mice was measured for 5 weeks post STZ treatment. Clinical signs of diabetes assessed as mice with blood glucose levels above 20mM (marked with grey dashed line). (c) Fluorescence microscopy images of pancreas sections taken from wild type and *Ppp1r15a^ΔC/ΔC^* mice fed either a normal chow diet (top panel) or 60% high-fat diet and streptozotocin treated (bottom panel). Fixed tissue sections were probed with antibodies against glucagon (green) and insulin (red), and counterstained with Hoescht (blue) and imaged by confocal microscopy. Scale bar = 50μm. (d) Circulating insulin levels of STZ-treated mice fed on a high-fat diet and shown compared to results from Figure 5c. (e) Glucose tolerance test (GTT) was carried out on 17-week, HFD+STZ treated mice. Mice were fasted overnight, and then injected with 1g/kg glucose. Blood glucose was measured at 0,10, 20, 30, 60, 90 and 120 minutes. N=7-9; two-way ANOVA and Bonferroni post-hoc test used for statistical analysis. (f) Quantification of the GTT carried out using the area under the curve (AUC). P value calculated by unpaired Student’s t-test. * p<0.05.

However, when challenged with an intraperitoneal injection of glucose, the *Ppp1r15a^ΔC/ΔC^* mice were significantly more glucose tolerant than wild type animals (Fig 6e). As expected, both genotypes had higher peak glucose levels compared with animals not treated with streptozotocin (compare Fig 5a & 6e), but despite having a partial depleted beta-cell mass, *Ppp1r15a^ΔC/ΔC^* mice no longer showed higher peak glucose levels compared to wild type controls, but still displayed an increased glucose clearance (Fig 6e).

## Discussion

We have shown that female mice lacking functional PPP1R15A eat less and gain less weight than wild type animals when fed a high-fat diet. This leads to reduced fat mass, improved insulin sensitivity, lower levels of circulating free fatty acids, and less hepatic steatosis in the *Ppp1r15a^ΔC/ΔC^* female mice.

Our observation that female *Ppp1r15a^ΔC/ΔC^* mice remained leaner than controls on a high-fat diet was unexpected. Despite reporting this allele more than a decade ago, we had never observed a weight phenotype in this line [12]. However, we had not previously challenged these animals with a high-fat diet. Likewise, the recent report that deletion of the *Ppp1r15a* gene generates male mice that spontaneously become obese and develop fatty liver was surprising to us [17]. In that study, *Ppp1r15a* deficient mice fed on standard chow (CLEA Rodent Diet CE-2) gained more weight than wild type controls, but this became significant only from 6 months of age – much older than is typical for our mice to be kept [17]. By contrast, a subsequent report by the same group failed to show excess obesity in female *Ppp1r15a* deficient mice even by 11-20 months of age [24]. Indeed, at 11-20 months of age on standard chow, 12 of 22 *Ppp1r15a* deficient females were described as being “lean”, compared with 0 of 37 wild type animals (Table 1 in reference [24]). This led us to wonder if female *Ppp1r15a* deficient mice might display a more subtle weight phenotype than had been reported for males. We therefore examined the effects of normal chow versus high-fat diets, which revealed reduced weight gain in *Ppp1r15a* deficient females only when fed a high-fat diet.

Differences between our results and those of Nishio and colleagues [17, 24] might also relate to the different alleles used. PPP1R15A is a large protein of >600 amino acids but only its C-terminal 150 residues are required for its phosphatase activity. We recently described how this C-terminal domain is both necessary and sufficient to interact with PP1 and actin to form a functional eIF2α holophosphatase [10, 29]. The different weight phenotypes between our *Ppp1r15a^ΔC/ΔC^* mouse versus the complete-deletion mouse may hint to the existence of, as yet, unidentified functions of the N-terminal portion of PPP1R15A. To test this, future strain-controlled studies will need to compare these two alleles in head-to-head experiments.

In keeping with the healthy phenotype of our *Ppp1r15a^ΔC/ΔC^* mice, no pathogenic mutations in *PPP1R15A* have yet been reported in humans. Importantly, mutation of *PPP1R15A* has not been identified in obese or lipodystrophic patients. By contrast, loss-of-function mutations of the constitutively expressed paralogue *PPP1R15B* have been shown to cause developmental defects both in mice [30] and humans [31]. Mice homozygous for an inactive mutant allele of *Ppp1r15b* survive gestation but then die during the neonatal period [30]. Recently, two siblings with early onset diabetes, short stature, microcephaly and learning disability were reported to have a homozygous mutation of the *PPP1R15B* gene [31]. The mutation affected a conserved arginine in the C-terminal PP1-binding domain of PPP1R15B and impaired formation of an active eIF2α holophosphatase when expressed *in vitro*. Although the mechanism by which PPP1R15B inactivation led to beta-cell dysfunction is unclear, it predicts that non-selective inhibition of P-eIF2α dephosphoryaltion might have negative developmental effects. This highlights the need to develop selective PPP1R15A inhibitors.

ER stress links obesity, lipotoxicity and the development of insulin resistance [32, 33]. Professional secretory cells, such as insulin secreting beta-cell, are prone to ER stress because they face large swings in the flux of new proteins traversing through their ER [34]. This in turn renders beta-cells especially vulnerable to defects in the UPR signaling pathway [5]. Indeed, mutations that impair the ability of mice or humans to phosphorylate eIF2α during ER stress have disproportionately greater effects on the beta-cell compared to other secretory tissues [4, 5, 14, 35, 36]. Because many solid tumours also depend upon eIF2α phosphorylation by PERK to withstand their hypoxic microenvironment [37], PERK inhibition has emerged as a potential chemotherapeutic strategy, but in pre-clinical models PERK inhibitors rapidly induce beta-cell toxicity, underling the reliance of this cell type on its ability to phosphorylate eIF2α [38, 39]. The precise mechanisms linking perturbed eIF2α phosphorylation with beta-cell toxicity are not fully understood, but recent observations suggest that proper levels of P-eIF2α are required to maintain appropriate levels of the chaperones required for proinsulin proteostasis [40].

Phosphorylation of eIF2α has complex effects on energy metabolism. When transgenic mice were generated to overexpress the C□terminal PP1 binding domain of PPP1R15A in the liver, they remained leaner than controls when fed a high-fat diet, and showed enhanced glucose tolerance, improved insulin sensitivity and reduced hepatic steatosis [25]. It is noteworthy, however, that phosphorylation of eIF2α had a strikingly biphasic effect on hepatic lipid metabolism with lipogenic gene expression being suppressed both at reduced and at elevated levels of P-eIF2α [25]. In addition, loss of hepatic P-eIF2α by overexpression of PPP1R15A has been shown to affect muscle and adipose insulin sensitivity, possibly through altered levels of circulating IGFBP-3 [41]. Our observations now indicate that P-eIF2α may also affect energy metabolism through effects on dietary intake.

Our observation that *Ppp1r15a^ΔC/ΔC^* mice fed a high-fat diet consume significantly fewer calories compared with controls is consistent with a central effect of elevating P-eIF2α. This is in keeping with the known role of eIF2α in regulating feeding behaviors. Aversion to diets imbalanced in amino acids has been shown to involve GCN2-mediated phosphorylation of eIF2α in the brain, while reduced eIF2α phosphorylation appeared to correlate with increased food intake [42]. Moreover, pharmacological enhancement of P-eIF2α in the mediobasal hypothalamus with salubrinal has been shown to reduce food intake [16]. Central ER stress has also been implicated in mediating hepatic effects in diet-induced obesity in mice, since overexpression of the chaperone Grp78 in the circumventricular subfornical organ reduced hepatic steatosis without affecting weight gain, food intake or adiposity [43]. However, we are unable to exclude an effect of *Ppp1r15a^ΔC/ΔC^* on the secretion of orexigenic or anorexigenic hormones that might regulate dietary choice.

Owing to the absence of a conditional knockout allele of *Ppp1r15a*, we were unable to determine in which tissues loss of PPP1R15A is responsible for the food intake phenotype. While this represents an important limitation of our study, the inactivation of PPP1R15A throughout the animal gives an indication of the effect PPP1R15A inhibition might have in a clinical setting. The complexity of the knockout phenotype, with better preservation of insulin sensitivity and yet higher peak glucose levels following glucose challenge, led us to question whether PPP1R15A antagonism would be beneficial or detrimental in models of diabetes. Our finding that *Ppp1r15a* mutant mice depleted of most beta-cells with streptozotocin have improved glucose tolerance suggests that the beneficial effects on insulin sensitivity dominate in the setting of diabetes. This raises the exciting possibility that PPP1R15A inhibition may be a potential therapeutic strategy in obesity with insulin resistance or in advanced states of the natural evolution towards diet-induced diabetes.

We previously showed that loss of PPP1R15A ameliorated ER stress-induced nephrotoxicity [12], which led us to speculate that inhibition of eIF2α might protect beta-cell mass in the context of beta-cell exhaustion. Indeed, deletion of *Chop*, a transcription factor responsible for efficient induction of PPP1R15A, was shown to protect mice in variety of models of diabetes [39, 44]. However, loss of *Ppp1r15a* has yet to be tested directly in an animal model of diabetes. Instead, efforts have focused on targeting PPP1R15A with small molecules, but these studies have been hampered by a lack of potent and selective molecules. Indeed, we recently demonstrated that the putative PPP1R15A inhibitors guanabenz and sephin1 do not affect PPP1R15A function, highlighting the urgent need for effective small molecules [45]. Salubrinal weakly inhibits dephosphorylation of eIF2α at high concentrations [45], but its dramatic effects to increase P-eIF2α in cells may reflect additional effects perhaps on one or more of the eIF2α kinases. In insulinoma cells, salubrinal was found to enhance the toxicity of palmitate [46], but our data indicate that genetic loss of PPP1R15A ameliorates the lipotoxicity of palmitate. This supports a potential therapeutic role of PPP1R15A inhibition if more potent and selective small molecules were to be generated.

In summary, we have observed that mice lacking the C-terminal PP1-binding domain of PPP1R15A are protected from obesity when fed a high-fat diet. This appears to reflect an increased satiety to diet, resulting in reduced weight gain and improved insulin sensitivity.

## Acknowledgements

We would like to thank Sarah Grocott and Dan Hart for technical support.

## Contribution statement

VP designed and performed the majority of experiments presented. SJM conceived the project as a whole. GB designed and helped to perform and anlyse experiments addressing the metabolic phenotype of the *Ppp1r15a* mutant mice. JEC, SC, AJTE, CG and LED contributed to experimental design and performance throughout. FMG, AVP and SJM oversaw the project. All authors contributed to writing the manuscript.

---

**Supplementary Figure 1. Inactivation of PPP1R15A reduces ER stress in response to tunicamycin** (a) Immunoblot for PPP1R15A, P-eIF2α, T-eIF2α, ATF4 and CHOP in lysates of *wild type* and *Ppp1r15a^ΔC/ΔC^* MEFs following treatment with tunicamycin (Tm) 2μg/mL for indicated times. Proteins of the expected sizes are marked with a solid triangle for PPP1R15A or an open triangle for PPP1R15A-ΔC.

(b) Quantification of (a) using ImageJ software.

(c-f) *Wild type* and *Ppp1r15a^ΔC/ΔC^* MEFs were treated with tunicamycin 2μg/mL for indicated times and RNA was prepared. *Ppp1r15a (Exons 1-3), Ppp1r15a (Exons 1-2), Atf4, and Chop* were quantified relative to *actb* by qRT-PCR. N=3; mean ± SEM. P value calculated by two-way ANOVA.

(g) Immnoblot for puromycin and tubulin in lysates of *wild type* or *Ppp1r15a^ΔC/ΔC^* MEFs following treatment with tunicamycin (Tm) 2μg/mL for indicated times. Ten minutes prior to harvesting, puromycin was added to the culture medium at a final concentration of 10ng/mL.

(h) Immunoreactivity to puromycin within lysates served as a marker of protein translation and was quantified using ImageJ software. *** p<0.001, ** p<0.01, * p<0.05.

